# Careful deployment of oilseed rape crops with *Rlm6* resistance gene against *L. maculans* is recommended to prevent the loss of efficacy of this resistance gene in French condiment mustard

**DOI:** 10.1101/297937

**Authors:** L. Bousset, M. Ermel, R. Delourme

## Abstract

Breeding varieties for increased disease resistance is a major means to control epidemics. However, the deployment of resistance genes through space and time drives the genetic composition of the pathogen population, with predictable changes in pathotype frequencies. In France, *Leptosphaeria maculans* causes disease on *Brassica napus* oilseed rape crops but not on *B. juncea* condiment mustard. Prior to the deployment of winter *B. napus* varieties with *Rlm6* resistance gene introduced from *B. juncea*, the aim of our study was to investigate if this deployment could impact disease control in condiment mustard. We assessed the presence of resistance genes against phoma stem canker in a set of current French *B. juncea* varieties and breeding lines. *Rlm6* was detected in all the 12 condiment mustard varieties. *Rlm5* was also detected in 8 varieties. No additional resistance genes were detected with the set of isolates used. Because frequency of isolates virulent on *Rlm6* is very low, these results indicate that *Rlm6* gene is a major component of disease control in the French *B. juncea* mustards tested. Using *Rlm6* in oilseed rape varieties will very likely induce an increase in frequency of *Rlm6* virulent isolates. This raises the acute concern of a wise deployment of oilseed rape around the condiment mustard growing area. Scientific knowledge on adaptation dynamics, spatial segregation of crops and cooperation between actors is currently available in order to mitigate the risk and advert negative consequences of the introduction of *Rlm6* resistance gene in oilseed rape varieties.

## Introduction

In crops, breeding varieties for increased disease resistance is a major means to control epidemics. The call to wisely manage the available host resistance genes is not recent, but still relevant. Even if focusing on resistance to one disease could inadvertently increase susceptibility to another (Arraiano & Brown, 2017), breeding for resistance to fungal pathogens contributed to yield increase (Potter et al., 2016).

However, the deployment of different resistance genes through space and time drives the genetic composition of the pathogen population (Hovmøller et al., 1997; Papaïx et al., 2011). Because epidemics of successive cropping seasons are not independent, the adaptation of pathogen populations to host resistances proceeds at the scale of a network of fields on which the selection pressures are not homogeneous, during a succession of cropping seasons (Bousset & Chèvre, 2013). Previous studies indicate that resistance genes in varieties exert directional selection for isolate-host compatibility (Hovmøller et al., 1993, Bousset et al. 2018). Predictable changes in pathotype frequencies were observed following commercial use of genes, or in field experiments (Hovmøller et al., 1993, Brun et al., 2010, Daverdin et al., 2012).

*Leptosphaeria maculans*, is the cause of stem canker on oilseed and vegetable brassicas, especially *B. napus*, *B. juncea* and *B. oleracea* (Mendes-Pereira et al., 2003). Epidemics are initiated in autumn, mainly by wind dispersed ascospores produced following sexual reproduction on stubble (Lô-Pelzer et al., 2009). Leaf spots are observed from autumn to early spring. Stem cankers develop from spring to summer, up to the time of harvest, due to the systemic growth of the fungal hyphae from leaf spots to the leaf petiole through vessels, and subsequently to the stem base (Travadon *et al*., 2009).

In *B. napus* there are more than 15 qualitative resistance genes conferring resistance to *L. maculans* (see Delourme et al., 2006; Rimmer, 2006; Elliott et al. 2015; Raman et al., 2013 for reviews). Their presence in host varieties can be detected using sets of *L. maculans* isolates to inoculate seedlings under controlled conditions. Phoma stem canker populations are diversified worldwide, with contrasted histories regarding the use of plant resistance genes (Rouxel et al., 2003a, Marcroft et al., 2012a, Zhang et al., 2016) and subsequent adaptation of pathogen populations (Balesdent et al. 2006; Liban et al., 2016, Van de Wouw et al. 2017). Populations of *L. maculans* collected from hosts with different resistance gene combinations harbor different infectivity allele frequencies (Van der Wouw et al., 2014, 2017).

When resistance genes have been deployed in oilseed rape varieties, phoma stem canker populations have repeatedly become adapted over a few years. Documented examples are adaptation to the *B. napus* genes *Rlm1* in Europe and Australia (Rouxel et al., 2003b); *LepR3* (*Rlm1*) and *LepR1* in Australia (Van de Wouw et al., 2014, 2017); *Rlm7* in Europe (Leflon, 2013; Winter et al., 2016); and *Rlm3* in Canada (Zhang et al., 2016). Thus, *L. maculans* has repeatedly evolved virulence against nearly all major resistance genes released in oilseed rape so far, sometimes with almost complete yield loss.

Thus, on *B. napus*, phoma stem canker is one of the many examples of loss of efficacy of genes following their use in varieties of this crop. However, a slightly different concern appears when the same fungus can cause epidemics on several related hosts, as are *B. juncea* and *B. napus*. In Australia, "juncea canola" varieties were commercialised in 2007 (Burton et al. 2008). Their resistance genes were unknown, but assumed to be different from those of the *B. napus* varieties (Marcroft et al. 2012b, Elliott et al. 2015). *Rlm5* and *Rlm6* are reported to be present in *B. juncea* (Chèvre et al. 1997; Barret et al. 1998; Balesdent et al. 2002, 2005), as well as a recessive resistance gene (*LMJR2*) (Elliott et al. 2015). Prior to the commercialization of "juncea canola" there were only small areas of *B. juncea* for condiment use, however, phoma stem canker infection was already occurring. Isolates virulent on *B. juncea* are widespread throughout the Australian canola growing regions (Elliott et al. 2015; Van de Wouw et al. 2017).

In France, *B. juncea* is not grown over wide acreages, however, in the Dijon area it has both a patrimonial importance and contributes significantly to the local economy, for the production of Dijon mustard. Formerly imported from Canada, mustard seed production has been significantly increased since 2010, up to 6000 ha in Bourgogne in 2016-2017 (http://www.cote-dor.chambagri.fr) and stable since (T. Guinet, personal communication). Phoma stem canker is not currently a disease problem on *B. juncea* (T. Guinet, personal communication). Intercrop *Brassica* species used to prevent nitrogen leaching are mainly *Sinapis alba*, or *Raphanus sativus* and only few are *B. juncea* selected for the biofumigating action of sinigrin. As opposed to Canada and Australia where virulent isolates have been detected in *L. maculans* populations (Liban et al. 2016, Van de Wouw et al. 2017), the frequency of isolates virulent on *Rlm6* is very low in France, and the one on *Rlm5* intermediate, locally as low as 43% (Rouxel et al. 2003b; Balesdent et al. 2005, 2006). Due to this, there is potentially only a very low frequency of isolates that can attack *B. juncea*, compared to *B. napus*. *Rlm6* gene has been introduced from *B. juncea* into winter *B. napus* (Chèvre et al. 2008). Prior to the deployment of such oilseed rape varieties in France, the aim of our study was to investigate to which extent this deployment could impact disease control in condiment mustard. We assessed the presence of most common genes against phoma stem canker, including *Rlm5* and *Rlm6*, in a set of the 12 current French *B. juncea* condiment mustard varieties and breeding lines.

## Materials and methods

### Fungus

A set of five differential isolates of *L. maculans* allowed detecting the presence of most genes frequent in French oilseed rape varieties. Isolates’ genotypes were known from previous publications (Table 1).

**Table 1.**
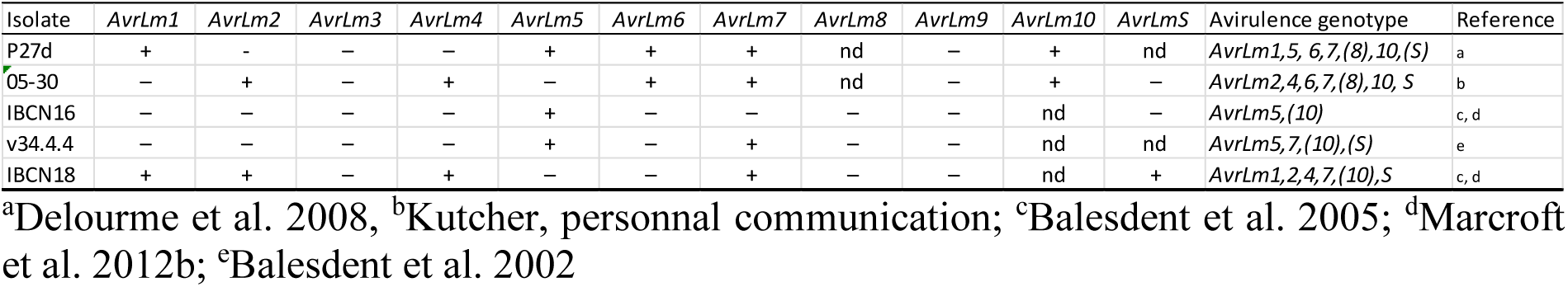
Presence/absence of the avirulence genes in the fungal isolate. Avirulence genotypes of *Leptosphaeria maculans* isolates based on disease scores on seedlings of Brassica lines. “+” refers to the presence of the avirulence allele; “-” refers to the presence of the virulence allele; “nd” refers to alleles that were not determined. The avirulence genotype indicates the avirulence loci for which the isolate has been characterized as avirulent; this nomenclature is as used by Balesdent et al. (2005).

Inoculum, consisting of suspensions of 10^7^ pycnidiospores per mL, was obtained for each isolate. Specifically, V8 agar (V8 juice 160 ml.l^−1^, agar 20 g.l^−1^) was autoclaved for 20 min at 120°C. After cooling, streptomycin concentrated solution 10% w/v in water was added to a final concentration of 0.1 g.l^−1^. For each isolate, agar plugs were transferred to V8 agar and grown for 10 days under near UV light. Pycnidiospores were dislodged from plates in sterile distilled water, filtered through muslin cloth. Spore concentrations were standardized to 10^7^ spores per ml after counting aliquots with a Malassez cell under a magnifying lens, aliquoted and stored at −20°C until tested for virulence.

### Plants and inoculation

After pregermination on wet filter paper, seeds of each variety were transplanted in trays with a 1:1:1 mix of sand, peat and compost and grown in climate chamber under a 16-h photoperiod for 10-12 days, at temperatures of 18°C night / 20°C day. A 10 µL drop of 10^7^ conidia per ml suspension was placed on each lobe of prick-wounded cotyledons (4 inoculation sites per plant) for four plants of each variety, in two independent trays. A plastic cover was placed over the inoculated plants to create a 100% relative humidity (RH) atmosphere for 24h in the dark. The three control lines ‘150-2-1’ (*Rlm5*, Balesdent et al. 2005), ‘Darmor’ (*Rlm9*, Delourme et al. 2004) and ‘DarmorMX’ (*Rlm6*, *Rlm9*) with known phenotypes were included in each test, so that both infective (virulent) and non-infective (avirulent) reactions were generated and the expression of interactions was confirmed.

### Scoring

Infection phenotypes were scored at 14dpi and confirmed at 21 dpi as described in Chèvre et al. (2008). The level of cotyledon infection was scored on a 0 to 11 scale in which 0 = no disease and 11 = cotyledons collapsed from phoma stem canker. The scale was modified from the 0 to 9 scale described earlier in which 0 = no darkening around wound – typical response of noninoculated controls or controls inoculated with water only; 1 = limited blackening around wound and lesion diameter of 0.5 to 1.5 mm; 3 = dark necrotic lesion with diameter of 1.5 to 3.0 mm; 5 = nonsporulating, 3 to 6 mm in diameter, lesion sharply delimited by darkened necrotic tissue; 7 = grayish green tissue collapse, lesion of 3 to 5 mm, with sharply delimited, nondarkening margin; and 9 = rapid tissue collapse at approximately 10 days, accompanied by profuse sporulation in large lesion (greater than 5 mm), and lesions with diffuse, nondarkened margins. To this scale an additional category of “11 = cotyledon totally collapsed, very often dry” was added (Chèvre et al., 2008) to categorize the most susceptible plants. Plants of classes 0 to 5 were considered as resistant as well as plants in classes 5 to 7 showing typical resistant plant reaction with blackening and limited symptoms that did not develop further between the two assessment dates at 14 and 21 days after inoculation. Plants of classes 9 to 11 were classified as susceptible, with symptoms appearing quickly, no blackening observed in lesions and with rapid disease progression between the two dates of assessment (14 and 21 days after inoculation). The scores were averaged over all infected plants.

## Results

### Expression of resistance and avirulence genes in control interactions

The susceptibility of ‘Darmor’ (lacking both *Rlm5* and *Rlm6*) to virulent isolates P27d, v34.4.4 and IBCN18 was confirmed (Figure 1). Isolates IBCN 16 and 05-30 gave slightly lower disease than expected.

**Figure 1.**
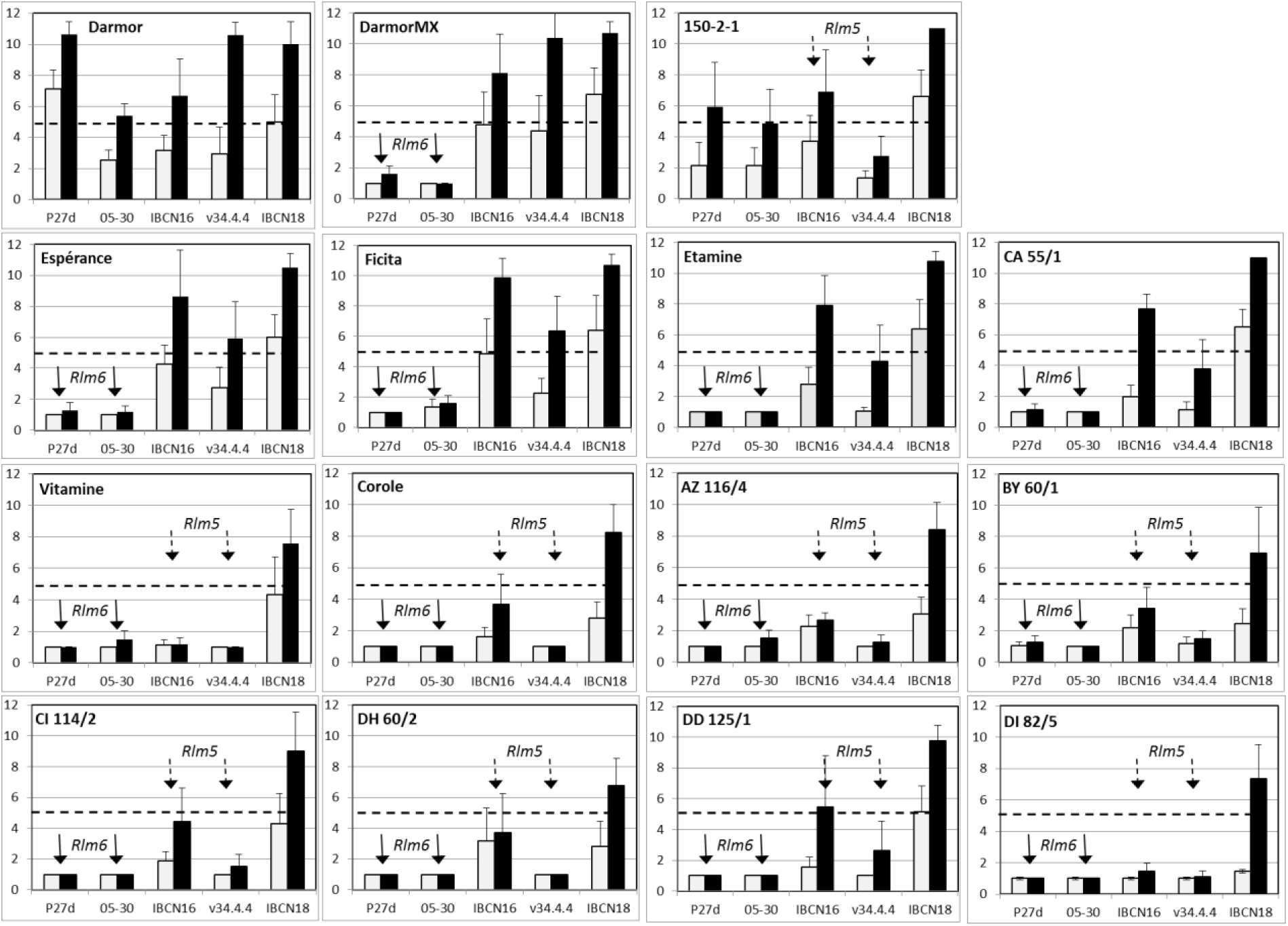
Mean disease scores at 14dpi (empty bars) and 21dpi (black bars) for the control lines ‘DarmorMX’ (*Rlm6*, *Rlm9*) ‘150-2-1’ (*Rlm5*) and ‘Darmor’ (*Rlm9*) and the 12 tested *B. juncea* mustards inoculated with *L. maculans* strains P27d, 05-30, IBCN16, v34.4.4 and IBCN18. Dashed line corresponds to score 5. Presence of *Rlm6* gene confers resistance to avirulent isolates P27d and 05-30 (Filled arrows). Presence of *Rlm5* gene confers resistance to avirulent isolates IBCN16 and v34.4.4 (Dashed arrows). Isolate’s profiles are given in Table1. Standard error is indicated‥.

The presence and expression of *Rlm6* gene in ‘DarmorMX’ was confirmed by the results at 21dpi with resistance against avirulent isolates P27d and 05-30 and susceptibility towards the virulent isolates v34.4.4, IBCN18 and IBCN 16.

The presence and expression of *Rlm5* gene in ‘150-2-1’ was confirmed by the results at 21dpi with resistance against avirulent isolate v34.4.4 and susceptibility towards the virulent isolates P27d, 05-30 and IBCN18. The isolate IBCN16 (*AvrLm5*) gave slightly more disease than expected on the ‘150-2-1’ control line, however interactions on mustards were clearly as with v34.4.4.

### Expression of resistance and avirulence genes in B.juncea condiment mustards

Presence of *Rlm6* gene was detected in all the 12 *B. juncea* condiment mustard varieties (Figure 1). *Rlm5* gene was also present in 8 varieties. The absence of *Rlm1*, *Rlm2*, *Rlm4*, *Rlm7* and *RlmS* in all the tested *B. juncea* was confirmed with the set of isolates used, and no additional resistance genes were detected. Resistance to isolates P27d and 05-30 and susceptibility towards the avirulent isolate IBCN 16 indicated the presence of *Rlm6* gene and the absence of *Rlm5* gene in the *B. juncea* condiment mustard lines Ficita, Espérance, Etamine and CA55/1 2 (Figure 1). Resistance to isolates IBCN16 and v34.4.4 as well as to isolates P27d and 05-30 indicated the presence of *Rlm5* and *Rlm6* genes in the *B. juncea* lines Vitamine, Corole, AZ116/4, BY60/1, CI114/2, DD125/1, DI82/5 and DH60/2 (Figure 1).

## Discussion

Only *Rlm6* and *Rlm5* genes were detected in the condiment mustard (Table 2). The absence of *Rlm1*, *Rlm2*, *Rlm4*, *Rlm7* and *RlmS* was confirmed with the set of isolates used, and no additional resistance genes were detected. Because isolates virulent on *Rlm6* are absent in French *L. maculans* populations (Balesdent et al. 2006), the results of our study indicate that *Rlm6* gene is a major component of disease control in the French *B. juncea* mustards tested. Breeding for resistance genes contributes to yield in canola varieties (Potter et al. 2016). *Rlm6* gene has been introduced into *B. napus* oilseed rape varieties (Chèvre et al. 2008) with the possibility of deploying these varieties in commercial production. The use of this gene in oilseed rape varieties will very likely induce and increase in frequency of *Rlm6* virulent isolates, as observed both in field experiments for this gene (Brun et al. 2010) and repeatedly in commercial situations for the other genes *Rlm1*, *Rlm3*, *Rlm7* (Rouxel et al. 2003b, Leflon, 2013, Van de Wouw et al., 2014, 2017, Winter et al. 2016, Zhang et al. 2016). Further, isolates virulent on *Rlm5* were already detected in French populations at frequencies locally up to 57% (Balesdent et al. 2006) weakening the disease protection conferred by this gene, and facilitating adaptation of the fungus. This raises the acute concern of a wise deployment of oilseed rape with *Rlm6* gene around the condiment mustard growing area.

**Table 2.**
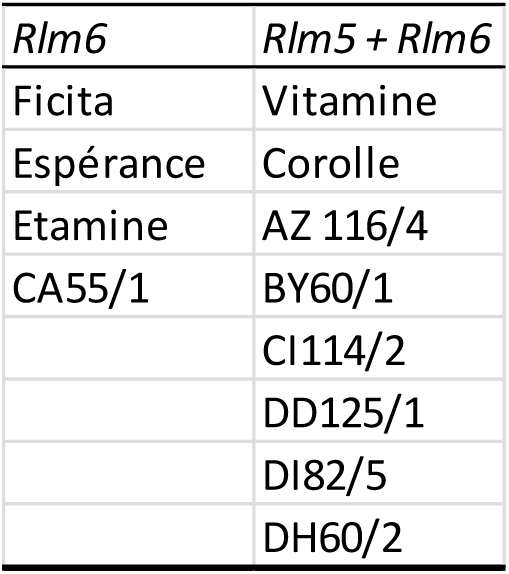
Presence of the seedling resistance genes in *B. juncea* condiment mustard. *Rlm1*, *Rlm2*, *Rlm4*, *Rlm7* and *RlmS* genes were absent from all varieties.

Combining qualitative and quantitative resistance within host lines has been shown to delay the loss of efficacy of qualitative resistance in a range of host-pathogen interactions, including fungi (Brun et al. 2010; Delourme et al. 2014), viruses and nematodes (Pilet-Nayel et al., 2017). The level of quantitative resistance in *B. juncea* is currently unknown. Because of the major consequences when qualitative resistance loose efficacy (Rouxel et al. 2003b, Van de Wouw 2014), an evaluation of the level of quantitative resistance in the *B. juncea* condiment mustards is desirable and methods are available.

While the call to wisely manage the available host resistance genes is not recent, it is still relevant. Importantly, the loss of efficacy of disease resistance genes can be averted when recommendation for deployment are produced in time (Van de Wouw et al. 2014). Local increase in disease severity on sentinel plots was detected for the variety Hyola50 in the Eyre peninsula, Australia. Recommendations of not sowing the varieties at risk in this area were followed, preserving their efficacy countrywide.

The deployment of different resistance genes through space and time drives the genetic composition of the pathogen population (Hovmøller et al., 1997; Papaïx et al., 2011). Modelling studies (Lô-Pelzer et al. 2010) and field experiments provide knowledge on the interplay between the acreages on which genes are deployed, the combination of genes in varieties, the intensity of inoculum sources and the spatial organization of crops in the landscape.

Without additional management strategies, the wider the acreage of the variety is, the faster and larger the increase in frequency of the corresponding virulence could be. There is currently no agreement on the best way to deploy available resistance genes, either individually or stacked in varieties (Rimbaud et al. 2018). On the one hand, diversifying selection has been proposed to play on pathogen’s weakness, such as stabilizing evolutionary dynamics (Zhan et al., 2015). On the contrary, stacking genes or QTLs in pyramids is also advocated (Pilet-Nayel et al. 2017). It is worth noticing that while stacking might be of interest when the corresponding virulences are absent from pathogen populations (Lof et al., 2017), this strategy leaves no possibility of leveraging the decrease of unnecessary virulences. In the case of the *B. napus* oilseed rape / *B. juncea* condiment mustard in France, using different gene pools is no longer possible because *Rlm6* gene is already present in *B. napus* varieties (Marcroft et al. 2012; Chèvre et al. 2008). Based on theoretical grounds in evolutionary biology of plant pathogen coevolutionary dynamics, it has been proposed that selectively reducing the contribution of pathogen populations from fields cropped with a resistant variety to the initial inoculum of the following season could slow down adaptation (Bousset & Chèvre, 2013). Field experiments with contrasting amounts of inoculum and contrasting levels of pre-adaptation (simulated by mixing stubble sources) confirm this hypothesis (Bousset et al., 2018). Thus, stabilizing the evolutionary dynamics should be considered, using options for reducing spore sources and reducing transmission by spatial organization of crops.

*L. maculans* survives on stubble. In field experiments, the reduction of inoculum by stubble management delayed adaptation (Daverdin et al. 2012). Currently, the previous year’s stubble is the primary contributing source for *L. maculans* (Marcroft et al., 2004) although changes in tillage cropping practice (McCredden et al., 2017) or shorter rotations might alter this situation. Options for reducing inoculum sources could include stubble management by burial (Marcroft et al., 2004), flooding or chemical application. Where possible, applying stubble management to reduce disease pressure on the mustards should be considered.

Wind dispersed ascospore produced by sexual reproduction are the main source of inoculum for *L. maculans* epidemics. Landscape structure is important for *L. maculans* transmission, such that increased isolation of crops (up to 500m) from any canola crops grown in the preceding year was associated with lower levels of disease (Marcroft et al., 2004). Transmission between fields can be predicted from spore dispersal (Marcroft et al., 2004; Bousset et al., 2015). Thus, spatially explicit models can be used to study and ultimately design combinations of landscapes, varietal choice and tillage practices promoting resistance durability against phoma stem canker (Lô-Pelzer et al., 2010; Rimbaud et al. 2018). As to the practicalities of spatial organization of the landscape, lessons have to be learned from the available case studies.

As to the optimization of the spatial segregation and the organization of actors, lessons can be learned from modelling studies on the segregation of GM-nonGM maize Bt crops in French regions Alsace, Sud-Ouest and Rhône-Alpes (Hannachi & Coléno, 2012). According to European recommendations, a farmer using GM seed has to use technical measures such as ensuring isolation distances to decrease cross-pollination risk and thus adventitious presence in the neighboring non-GM fields. For grain merchants, the problem is to ensure segregation of the two products in their supply chain and, as far as possible, to manage cross-pollination risk between GM and non-GM fields in their collecting area (Coléno et al. 2009). Admixture of products in the supply chain is not relevant for the *B.juncea* / *B.napus* case, but cross-pollination issues have similarities with *L. maculans* spore flows between fields. A model of farmer’s choice between GM and non-GM varieties for each field in a zone (Coléno, 2008) was combined with a spatially-explicit model of gene flow, MAPOD^®^ (Angevin et al., 2008) including pollen production, dispersal and pollination given distances, flowering times and varieties, to assess the consequences of crop placement (Coléno et al. 2009). The presence of GM in non-GM batches as a result of cross-pollination depends on the size of the non-GM zone and prevailing wind. Further modelling was used to compare strategies of either spatial or temporal segregation of infrastructure use for contrasting percentages of GM maize (Hannachi & Coléno. 2015). In all cases, cooperation between the four collection and storage agents proved better than competition. As spore dispersal is documented in *L. maculans* (Bousset et al. 2015), such an approach could be developed to understand the consequences of the placement of oilseed and mustard crops on the adaptive dynamics of the pathogen population. Alternately, such modelling could be a useful means to promote discussions between actors (Hossard et al. 2013).

The oilseed rape production is not currently organizing the spatial deployment of resistance genes for disease protection (Hannachi et al. 2017). However, for at least two oil quality-related issues, it is more than 10 years that French oilseed rape producers do organize spatial patterns by combining varietal choice and spatial segregation (Hannachi et al. 2017). First case is “Fleur de colza” sector to ensure Omega 3 contents and oil quality, ensuring the cooperation between one industrial actor, seven collection and storage agents and more than 1000 contracting growers in regions Centre, Ile-de-France et Bourgogne. A charter allows combining varietal choice and spatial deployment in the landscape. Second case is the « Pollen » collective organization to manage the cross-pollination in oilseed rape between low-erucic varieties (oil for human consumption) and high-erucic varieties (industry) suspected of risk in case of human consumption. By contracts and bonus, this collaboration involving breeders, industries, transfer and 3000 growers, allows to organize the landscape over 20 000 ha, for varietal choice, isolation distances and temporal rotations. The specifics of these two cases are the collective creation of additional value (higher incomes) inducing actors to cooperate, and mitigating the extra costs. Noteworthy, these additional values and common interests were not pre-existing, but derive from the collective cooperation (Hannachi et al. 2017). Dijon condiment mustard is a patrimonial product, locally anchored, with a worldwide reputation. The conditions under which cooperation with oilseed rape production could be established deserve further studies.

In conclusion, this study detects a threat for *L. maculans* disease control on French condiment mustard. Scientific knowledge is currently available in order to mitigate the risk and advert negative consequences of the introduction of *Rlm6* resistance gene in oilseed rape varieties. We thus recommend that around the condiment mustard growing area, only a good level of quantitative resistance or resistance genes different from *Rlm6*, are used in oilseed rape varieties for the control of *L. maculans*.

## Acknowledgements

We thank Claude Domin for technical assistance. We thank Thierry Guinet (AgroSup Dijon) for providing the condiment mustard seeds and the BraCySol Biological Ressource Center for providing oilseed rape seeds. We thank Mylène Balesdent for providing the *L. maculans* v34.4.4 isolate and Randy Kutcher for providing the *L. maculans* 05-30 isolate. We thank Mourad Hannachi for fruitful discussions. This work benefited from the financial support of INRA – the French National Institute for Agronomical Research. The authors declare the absence of conflict of interest. ME carried out experiments, LB and RD conceived and designed the study and prepared the manuscript, read and approved by all authors.

